# How the Motor Aspect of Speaking Influences the Blink Rate

**DOI:** 10.1101/2020.07.31.230391

**Authors:** Mareike Brych, Supriya Murali, Barbara Händel

## Abstract

The blink rate increases if a person indulges in a conversation compared to quiet rest. Since various factors were suggested to explain this increase, the present study tested the influence of motor activity, cognitive processes and auditory input on the blink rate but at the same time excluding any social interaction. While the cognitive and auditory factors only showed a minor influence, mere mouth movements during speaking highly increased the blink rate. Even more specific, lip movements, but less jaw movements, are likely responsible for the increase during a conversation. Such purely motor related influences on the blink rate advise caution when using blinks as neurological indicators during patient interviews.

## Introduction

Blink rate is assumed to reflect perceptual or cognitive load [1, 2]. Also, the often found increase in blink rate during conversation has been proposed to reflect cognitive processes besides several other aspects that might be influential. Doughty [3] summarizes a variety of internal states that can influence the blink rate, such as engagement, emotions or opinions. Hömke, Holler and Levinson [4] further showed that blinks can also be communicative signals between conversation partners. Other research has shown that the motor act of speaking [5], but not mere jaw movements produced during gum chewing [2] or the mere act of keeping the mouth open [6] increased blinking. Contradictory to the main results in the latter study, a small group that exhibited notable mouth and jaw movements during a no-task condition nearly had a doubled blink rate compared to those who did not show such movements. These inconsistencies are worrisome since blinks serve as neurological indicators in clinical settings. For example, Parkinson’s disease is associated with very low blink rates [7], while high blink rates are observed in patients with Schizophrenia [8]. What causes these deviations from normal blink behavior is not known. We set up an experiment to systematically investigate influences of facial motor activity on our blinking behavior, while at the same time we control for cognitive and auditory influences.

Considering human facial anatomy, it is known that the facial nerve (7^th^ cranial nerve) innervates the muscles for facial expressions and eyelid closing, but is not directly involved in chewing movements. These muscles are innervated by the trigeminal nerve (5^th^ cranial nerve) [9]. Whenever the facial nerve is malfunctioning, blinking is ceased and the corner of the mouth drops on the affected side [10]. During surgeries, facial nerve stimulation is also used to predict the postoperative function by checking motor-evoked potential in the eye ring muscle (orbicularis oculi) and the kissing muscle (orbicularis oris) [11]. This would predict that lip movements and blinking are closely coupled while chewing movements might be less connected to blinking. This could explain the negative result by Karson [2], who showed no relationship between chewing and blink rate, but would predict an increase in blink rate due to lip movement. Based on the anatomical findings we therefore hypothesize that the movement during talking accounts strongly for the blink increase during a conversation. We further expect the largest effect due to lip movement and not so much jaw movements, which are not so closely connected to the eyelid.

In order to account for the auditory and cognitive aspect of speaking, we included condition during which participants had to engage in an internal verbal discourse, which we call “talk inside the head”, or listen to a replay of their own talking. However, since hearing ourselves talk is rather unusual, we added a condition in which participants listened to someone else. Adding to our hypothesis that blink rate is mainly increased by motor related factors, we expected only a minor influence of auditory input or cognitive aspect of speaking on blinking. Visual stimulation as well as social influence were minimized in our experiment. Under this condition, our results confirmed a major influence of motor activity especially of the lips on blinking, while cognitive and auditory aspects only showed a minor influence.

## Methods

### Participants

30 psychology students of the University of Würzburg (mean age: 20.17 years ± 1.86 SD, 2 male) took part in the study. All participants gave their written informed consent and received study credit for their participation. The study was approved by the local ethics committee and was in line with the European general data protection regulations (DSVGO).

### Procedure

Participants sat alone in a noise shielded, very small, dimly lit room. They were allowed to freely move their eyes and head. Auditory instructions were given by a Sennheiser PC3 Chat headset. Binocular eye movements were recorded with 120Hz using SMI eye tracking glasses.

The study consisted of 8 conditions, which were repeated 5 times (except for the baseline which was repeated 15 times) and each lasted for 1 minute. During the baseline condition, participants had no task. During “normal talking“, “talking inside the head“ and “talking without sound“, participants were instructed to talk about easy topics like “Describe your apartment“ or “Describe your last holiday“. “Talking inside the head“ involved no mouth movement, while „talking without sound“ referred to simply mouthing words. To induce lip movements independent of talking, participants were asked to suck on a real lollipop (“lip movement“). In another condition (“jaw movement“), chewing a gum resulted in jaw movements. In the auditory conditions, auditory input was displayed by either a monologue of a young woman (“listen to someone else“) or their own monologue of a previous “normal talking“ trial (“listen to oneself“). The order of conditions was randomized, except that the condition “listen to oneself“ needed to be placed accordingly after the “normal talking“ condition. Participants were able to start each trial by pressing a button followed by a starting tone. The end of the trials was signaled by another tone.

### Data analysis

Four participants were excluded (three due to more than 20% eye data loss, one due to an extremely high mean blink rate >50 blinks/min). Additionally, the eye recording of one participants was lacking two trials, which resulted in a list-wise exclusion for some parts of the analysis. Recorded speech was digitally transformed into waveforms, which were controlled for outburst signaling continuous talking.

The experimental program was implemented and analyzed in MATLAB R2015b (Mathworks). Bayesian analysis was performed with JASP (JASP Team (2019), Version 0.11.1.0).

### Blink detection

We developed a blink detection algorithm based on pupil radius. Blinks were initially detected when both z-transformed radii were below a threshold of −2 standard deviations. The start and the end of the blink were then shifted to the time point when the radii were higher than half the threshold. Blinks less than 100ms apart from each other were concatenated. Blinks longer than 1000ms and shorter than 50ms were discarded.

## Results

To test for cognitive influences on the blink rate, we compared baseline (no task) with “normal talking” and with “talking inside the head”. A repeated measures 2-factor ANOVA compared the blink rate between these conditions as well as between the five repetitions of each condition within subjects. A significant main effect of conditions was revealed (*F*(1.31, 32.86) = 25.22, *p* < .001, η_p_^2^ = 0.50, Greenhouse-Geisser corrected (GG)). Post-hoc pairwise t-tests revealed a significant difference between normal talking and talking inside the head (*p* < .001) as well as between baseline and normal talking (*p* < .001). There was neither a significant main effect of repetition nor a significant interaction effect (both F < 1) (Fig.1a).

**Figure 1.**
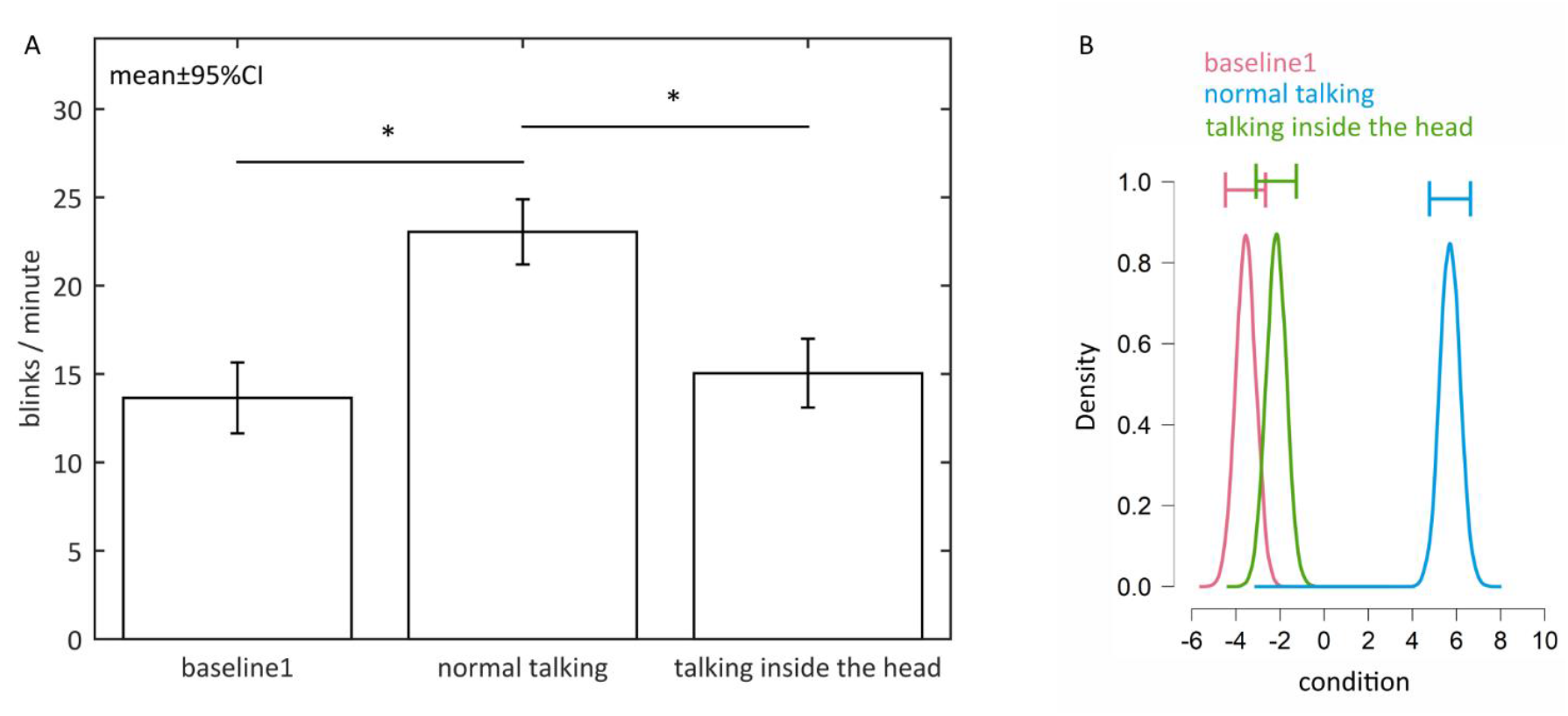
Effects of no task, normal talking and “talking inside the head” on the blink rate. A. Blink rate during the baseline condition (no task), normal talking and “talking inside the head”. Error bars represent 95% confidence intervals. Stars mark significant differences revealed by parametric statistics. B. Posterior distributions of the effect of each condition on the blink rate. Normal talking has highest effect on blink rate followed by “talking inside the head” and baseline. The horizontal error bars above each density represent 95% credible intervals.

In order to assess the magnitude of differences between conditions, we additionally computed a Bayesian analysis. Bayesian ANOVA similarly revealed overwhelming evidence that the conditions had a very robust effect on the blink rate (Bayes Factor: BF_10_ = 7.033*10^26^). Post hoc tests showed evidence that blink rate during “normal talking” differed to blink rate during baseline as well as to blink rate during “talking inside the head” (adjusted posterior odds of 2.606*10^16^ and 1.686*10^11^). Additionally, there was small evidence that baseline and “talking inside the head” were the same (adjusted posterior odds of 1.212) (Fig. 1b).

Comparing the blink rate between motor components revealed a high blink rate during “talking without sound”, followed by lip movements and jaw movements. The condition with no movement showed the lowest blink rate. A repeated measures 2-factor ANOVA showed a significant main effect of these conditions on blink rate (*F*(2.19,52.57) = 9.00, *p* < 0.001, η_p_^2^ = 0.27 (GG)). Post-hoc pairwise t-test specified this effect. The blink rate was significantly lower during the baseline condition compared to lip movements (*p* = .023) and compared to “talking without sound” (*p* = .002). Again, we found neither a main effect of repetition (*F*(2.13,51.21) = 2.46, *p* = .917, η_p_^2^ = .09 (GG)) nor an interaction effect (*F*(4.20,100.77) = 1.15, *p* = .338, η_p_^2^ = .05 (GG)). The difference between jaw movement and baseline did not reach significance (Fig.2a).

**Figure 2.**
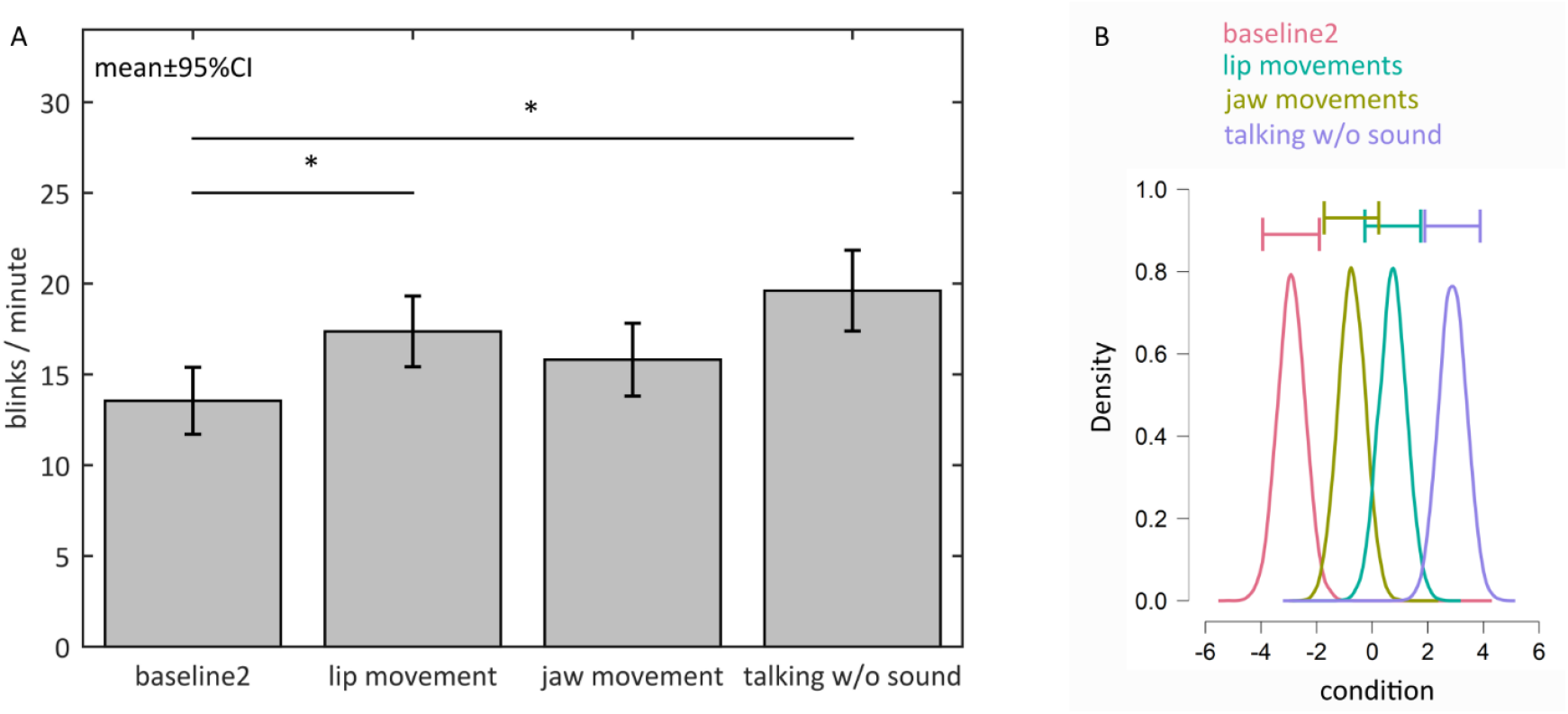
Effects of no task, jaw movements, lip movements and “talking without sound” on the blink rate. A. Blink rate during the second baseline condition (no task), moving the lips during lollipop sucking, moving jaw muscles during gum chewing and talking without sound production. Error bars represent 95% confidence intervals. Stars mark significant differences revealed by parametrical statistics. B. Posterior distributions of the effect of each condition on the blink rate. Talking without sound has highest effect on blink rate followed by lip movement, jaw movement and baseline. The horizontal error bars above each density represent 95% credible intervals.

Again, Bayesian ANOVA supported the effect of conditions on blink rate (Bayes Factor: BF_10_ = 4.749*10^8^). Post-hoc comparisons revealed strong evidence for differences in blink rate between baseline and lip movements as well as between baseline and “talking without sound” (adjusted posterior odds of 2.267*10^3^ and 5.018*10^5^). There was also evidence for differences in blink rate between baseline and jaw movements as well as between jaw movements and “talking without sound” (i.e. odds of 15.654 and 32.688). Blink rates during lip movements and jaw movements as well as during lip movements and “talking without sound” were similar (i.e. odds of 1.337 and 1.661) (Fig.2b).

Additionally, a significant difference in blink rate when comparing the baseline condition with listen to someone else and listen to oneself was found (F(1.48,35.61) = 3.74, p = .045, η_p_^2^ = 0.13 (GG). Post-hoc tests however did not reveal a difference between the baseline condition and any auditory input (*p*s > .116), but a significant difference between listen to oneself and listen to someone else (*p* = .020). Following the analysis for the previous questions, we found neither a significant main effect of repetition (F(2.78, 66.73) = 1.91, p = .141, η_p_^2^ = .07 (GG)) nor a significant interaction effect (F(4.25,102.02) = 2.04, p = .091, η_p_^2^ = .08 (GG)) (Fig.3a).

**Figure 3.**
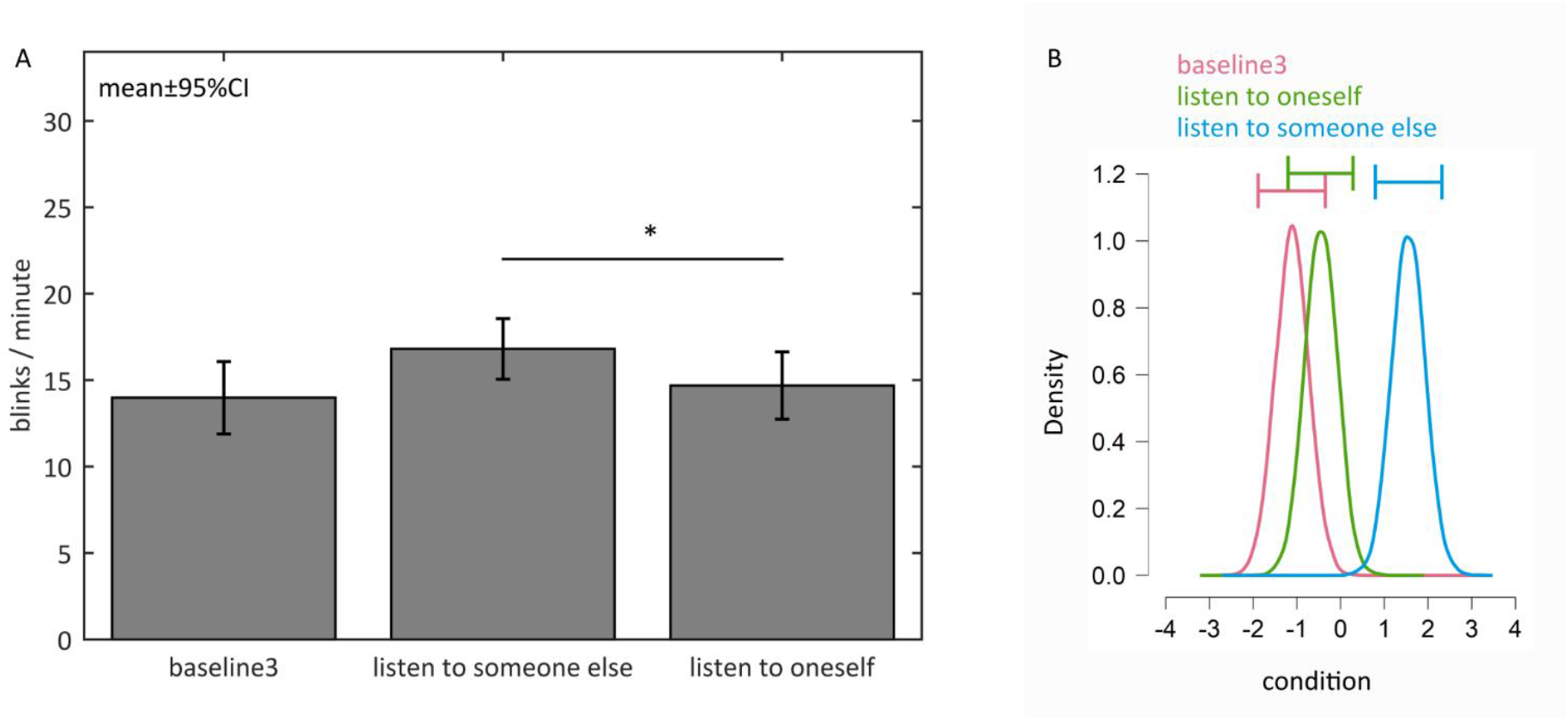
Effects of no task, listen to someone else and listen to oneself on the blink rate. A. Blink rate during the third baseline condition (no task), listening to someone else and listening to a previously recorded monologue. Error bars represent 95% confidence intervals. Stars mark significant differences revealed by parametric statistics. B. Posterior distributions of the effect of each condition on the blink rate. Listening to someone else has highest effect on blink rate followed by listening to oneself and baseline. The horizontal error bars above each density represent 95% credible intervals.

While the Bayesian ANOVA again revealed a strong effect of conditions (Bayes Factor: BF_10_ = 183.819), post-hoc tests showed a slightly different picture. There was slight evidence for a difference in blink rate between baseline and listen to oneself (adjusted posterior odds of 1/0.097 = 10.309), as well as between listen to oneself and listen to someone else (i.e. odds of 10.646) and stronger evidence for a difference in blink rates between baseline and listen to someone else (i.e. odds of 47.914) (Fig.3b).

## Discussion

Our results replicated previous findings that talking is accompanied by an increase in blink rate compared to baseline [e.g. 2]. More specifically, our findings enable to identify that neither the cognitive processes nor the auditory input, but rather, the motor activity of the mouth has the main influence on our blink rate.

The conditions “talking inside the head” and “normal talking” differ in terms of motor output and auditory input but not cognitive processes, which are needed for the production of meaningful sentences. Since the blink rate is significantly lower during “talking inside the head” than during normal talking and highly similar to the baseline, cognitive processes during speaking seem to have, if at all, little effect on our blinking. Whether cognitive influences during a real conversation might have an effect on blinking, cannot be excluded with our setup.

Also, the self-induced auditory input due to talking is not the cause of the increase in blink rate during talking since the blink rate during normal talking is only slightly higher than during talking without sound (23.05±1.84 compared to 19.62±2.22 (mean±95%CI)). However, Bayesian analysis showed that there is at least some evidence that auditory input influences the blink rate. Nevertheless, that listening to someone else showed a higher blink rate than listening to oneself suggests additional influences. In contrast to our findings, Bailly, Raidt and Elisei [12] further suggest an inhibition of blinking during listening periods within a conversation compared to waiting periods. There are some possible reasons for this difference, one lying in the difference of the setup. While during a conversation one is bound to attend to the auditory input of the conversation partner in order to respond accordingly, in our experiment, the auditory input was non-task relevant. However, the differences might also be explained by the fact that our experiment explicitly excluded social interaction. Having a conversation with a real partner might change our blinking behavior. Indeed, it was shown that the duration of blinks can serve as a feedback signal for the conversation partner [4] and that speakers often blink at the end or during pauses in speech [13].

Overall, our findings that talking without sound as well as lip movements during lollipop sucking increased blinking clearly suggests that motor related influence are the main cause for increased blink rate while talking, at least in a situation outside a conversation with a partner. More specifically, by separately investigating the influence of different muscle groups, our results indicate that some of them are more strongly linked to blinks than others. Chewing movements are not sufficient to significantly increase the blink rate when using a parametric statistical approach, a finding that is in line with previous research [2]. However, our Bayesian analysis still suggests a weak link. The lip movements on the other hand show a clear effect on blink rate. These muscles as well as the eye muscles are innervated by the facial nerve and might be activated together, while somewhat further connections are not so closely coupled [9]. Apart from close neuronal connectedness, previous research revealed various interactions between movements that are not based on close-by nerves. This suggests that there also might be a common phenomenon of motor interaction. For example, finger tapping entrains spontaneous blinking [14] as does walking speed [15]. Similarly, other eye movements like (micro-)saccades co-occur with head movements [16, 17] and a large saccade size comes with a high blink probability [1]. Nissens and Fiehler [18] could also show that saccades and reach movements can influence each other’s trajectories.

In addition, the magnitude of blink enhancement might be related to the amount of muscles that are involved in articulation, which is varied by the frequency or complexity of the motor activity. During a conversation, we normally speak at a rate of 5.3 syllables per second [19], which refers to approximately 200 words per minute (language dependent). Although it seems plausible that talking has the highest movement rate compared to lollipop sucking and gum chewing, a detailed analysis needs to be performed to assess any relationship. The influence of articulation complexity on blinking was touched when comparing the possibly more complex mouth movements during reciting numbers from 100 upward and the simpler movements during reciting the alphabet [5]. The authors concluded that more complex movements intensified our blinking.

In summary, we showed that the motor activity during speaking has a major influence on blinking, while auditory input and cognitive processes only have a minor effect. Given these results, we advise caution when using blinks as neurological indicators during patient interviews without closely monitoring the time of talking.

